# Evidence for a long-range RNA-RNA interaction between *ORF8* and *Spike* of SARS-CoV-2

**DOI:** 10.1101/2021.11.09.467911

**Authors:** Okiemute B. Omoru, Filipe Pereira, Sarath Chandra Janga, Amirhossein Manzourolajdad

**Author notes:** Correspondence should be addressed to: Amirhossein Manzourolajdad, Informatics and Communications Technology Complex, 535 West Michigan Street, Indianapolis, IN 46202.

## Abstract

SARS-CoV-2 has affected people worldwide as the causative agent of COVID-19. The virus is related to the highly lethal SARS-CoV responsible for the 2002-2003 SARS outbreak in Asia. Research is ongoing to understand why both viruses have different spreading capacities and mortality rates. Like other beta coronaviruses, RNA-RNA interactions occur between different parts of the viral genomic RNA, resulting in discontinuous transcription and production of various sub-genomic RNAs. These sub-genomic RNAs are then translated into other viral proteins. In this work, we performed a comparative analysis for novel long-range RNA-RNA interactions that may involve the *Spike* region. Comparing predictions between reference sequences of SARS-CoV-1 and SARS-CoV-2 revealed several predictions amongst which a thermodynamically stable long-range RNA-RNA interaction between (23660-23703 *Spike*) and (28025-28060 *ORF8*) unique to SARS-CoV-2 was observed. Using data gathered worldwide, sequence variation patterns observed in the population support the *in-silico* RNA-RNA base-pairing predictions within these regions, suggesting further evidence for the interaction. The predicted interactions can potentially be related to the regulation of sub-genomic RNA production rates in SARS-CoV-2 and their subsequent accessibility to the host transcriptome.

## Introduction

Severe acute respiratory syndrome coronavirus 2 (SARS-CoV-2) is a highly transmissible and pathogenic coronavirus that emerged in late 2019 and has caused a pandemic of acute respiratory disease, named coronavirus disease 2019 (COVID-19) (Hu, Guo et al. 2020). SARS-CoV-2 is related to SARS-CoV (denoted here as SARS-CoV-1), a life-threatening virus responsible for an outbreak in 2002-2003 that was contained after intense public health mitigation measures (Standl, Jockel et al. 2020). The Coronaviruses belong to the *Coronaviridae* family. They are enveloped, positive-sensed, and have a single-stranded RNA genome (Cui, Li et al. 2019) and are categorized into different genera based on their protein sequences (Phan, Ngo Tri et al. 2018). While certain genera are non-patheogneic in Humans (Su, Wong et al. 2016, Wong, Lui et al. 2016), the genera of beta-coronaviruses comprise most human coronaviruses (HCoVs), including the SARS-CoV-1, MERS-CoV, HCoVOC43, HCoV-HKU1, and SARS-CoV-2 (Malik 2020). Beta-coronaviruses, including SARS-CoV-2, are highly pathogenic and are responsible for life-threatening respiratory infections in humans.

The SARS-CoV-2 genome is approximately 30’000 nucleotides long. The nucleotide content of the viral genome consists majorly of two large open reading frames (ORF1a and ORF1b) and structural proteins (Su, Wong et al. 2016, Chen, Liu et al. 2020). The two ORFs translate into replicase polypeptides (pp1a and pp1b) which are digested by viral proteases to produce the 16 non-structural proteins (NSPs 1-16) that aid in viral replication and transcription (Khailany, Safdar et al. 2020, Naqvi, Fatima et al. 2020, Rahimi, Mirzazadeh et al. 2021). The sub-genomic RNAs encode for the four structural proteins, spike (S), envelope (E), membrane (M), and nucleocapsid (N) proteins, as well as several accessory proteins known as Open Reading Frames (ORF) 3a, 6, 7a, 7b, 8, and 10. The structural proteins are responsible for viral assembly and suppressing the host’s immune response (Kim, Lee et al. 2020, Yao, Song et al. 2020). SARS-CoV-2’s viral proteins -NSPs and structural proteins also play an important role in the viral entry, life cycle, and overall survival of the virus in the host cell.

The first steps of coronavirus infection involve the viral entry into the host cell via binding of the Spike (S) protein to the cellular entry receptors for attachment to the receptor-binding site of the hosts cell membrane, fusion, and the release of the viral RNA into the cell. In humans, the host cellular receptor for SARS-CoV-2 is human angiotensin-converting enzyme 2 (ACE2) (V’Kovski, Kratzel et al. 2021). The interaction between Spike and ACE2 determines the viral response and pathogenicity. (Li 2016, Tortorici and Veesler 2019, Letko, Marzi et al. 2020). After entry, SARS-CoV-2 expresses and replicates its genomic RNA to produce full-length copies, integrated into the newly created viral particles (V’Kovski, Kratzel et al. 2021). SARS-CoV-2’s genome encodes NSPs, which are essential for viral RNA synthesis, and structural proteins necessary for virion assembly (Shin, Jung et al. 2018).

Coronavirus RNA-dependent RNA synthesis includes two differentiated processes of genome replication and transcription of a collection of sub-genomic RNAs. The sub-genomic RNAs encode the viral structural and accessory proteins. These RNAs are produced by discontinuous transcription where the synthesis of the negative-sense strand is disrupted. The resulting strand will then produce a plus RNA strand sub-genomic RNA. The complex replication/transcription machinery production of a series of sub-genomic RNAs through the process of template switching during negative-sense RNA synthesis (Sola, Almazán et al. 2015, Snijder, Limpens et al. 2020). The replication of the viral structural protein RNAs is mediated by RNA-dependent RNA polymerase (RdRp). The S, E, and M proteins are translated by ribosomes attached to the endoplasmic reticulum. N protein remains in the cytoplasm and is assembled from genomic RNA. It then fuses with the virion precursor and is transported from the endoplasmic reticulum and released to the cell surface via exocytosis (Fehr and Perlman 2015, Scudellari 2021).

Beta-coronaviruses can form long-range high-order RNA-RNA interactions that contribute to template switch and consequently regulate the viral transcription and regulatory pathways for the production of sub-genomic mRNAs (Sola, Almazán et al. 2015). Long-range interactions are generally found in positive-strand viruses (Nicholson and White 2014, Chkuaseli and White 2018). The longest RNA-RNA interaction found so far is ∼26000 and is involved in a sub-genomic RNA synthesis in coronaviruses (Nicholson and White 2014). Mediated by stabilizing proteins, such interactions impact the tertiary structure of the genomic RNA, facilitating binding of the 5’ UTR Transcript Regulatory Sequences (TRS) to the regulatory sequence upstream of a particular gene, leading to the switching of minus strand template to that of the gene’s sub-genomic transcript. Regulation of the *N*-gene sub-genomic transcript is a fair example of such high-order RNA-RNA interactions (Sola, Almazán et al. 2015). Although some efforts have been made to investigate RNA-RNA interactions in in general of SARS-CoV-2 (Ziv, Price et al. 2020), It is very difficult to identify all the genomic RNA regions that are involved in such intricate interactions, presenting challenges to finding novel interacting regions within the virus (Nicholson and White 2014).

The co-evolution of coronaviruses with their hosts is navigated by genetic variations made possible by its large genome (Woo, Huang et al. 2010), recombination frequency (of up to 25% for the entire genome in vivo) (Baric, Fu et al. 1990, Singh, Pandit et al. 2021), and a high mutation rate (Sánchez, Gebauer et al. 1992, Vijgen, Lemey et al. 2005). SARS-CoV-2’s mutation occurs spontaneously during replication. Thousands of aggregate mutations have occurred since the emergence of the virus (Li, Wang et al. 2020). A significant cause of concern about SARS-CoV-2’s mutations is a change that could lead to a highly lethal infection or a failure on the effects of the current vaccines (Collier, De Marco et al. 2021). It is known that the strain with the highest similarity to SARS-CoV-2 is SARS-CoV-1. Similar to SARS-CoV-2, SARS-CoV-1 has a genome length of around 30kb (29’751 nt), and its similarity ratio to the SARS-CoV-2 genome is 82.45% (Chen, Boon et al. 2021). The genomic differences explain the disparities in both viruses’ dispersal and immune evasion (Ortiz-Fernandez and Sawalha 2020). The percentage similarity of the Spike protein of SARS-CoV-2 and SARS-CoV-1 is 97.71%. Spike protein’s Receptor Binding Domain (RBD) which is the most variable part of the coronavirus genome (Tai, He et al. 2020), has 74.41% similarity. In fact, computational analysis has affirmed that the RBD sequence of SARS-CoV-2 differs from those observed to be ideal in SARS-CoV-1 (Wan, Shang et al. 2020); hence, the high-affinity binding of the SARS-CoV-2 RBD to the human ACE2 is consequently due to natural selection on human ACE2, which allows for a solution for binding (Walls, Park et al. 2020). A significant difference between the Spike regions of both viruses is a polybasic insertion at the S1/S2 cleavage site, resulting from a 12-nt insert in the Spike region of SARS-CoV-2 that does not exist in SARS-CoV. In addition to increasing Spike protein infectivity, the 12-nt insert may also have a role on the RNA level, since it has a high GC content (CCUCGGCGGGCA; positions 23,603-23,614 of the reference). Similarity of other structural proteins are as follows: E-96%, M-89.41%, and N-85.41%. The similarity between the structural protein of SARS-CoV-2 and other Coronaviruses is less than 50% (Cosar, Karagulleoglu et al. 2021).

RNA structures can play critical roles in the life cycle of Beta-coronaviruses. For instance, studies have reported that SARS-CoV-2’s genomic RNA occupy some of the hosts MiRNAs that control immune regulated genes, thus depriving them of their function (Zhang, Amahong et al. 2021). Recent studies have found locally stable RNA structures within the SARS-CoV-2 genome (Andrews, Peterson et al. 2020, Bartas, Brázda et al. 2020, Lan, Allan et al. 2020, Simmonds 2020). Moreover, *in-vivo* RNA structure prediction methods such as dimethyl sulfate mutational profiling with sequencing (DMS-MaP-seq) suggest that SARS-CoV-2 forms RNA structures within most of its genome (Lan, Allan et al. 2020), some of the possible relevance to the virus life cycle. These RNA structures can potentially be the target of RNA-based therapeutic applications (Manfredonia, Nithin et al. 2020, Rouskin, Lan et al. 2021), or may lead to methods for inhibiting viral growth (Huston, Wan et al. 2021).

The Spike gene has been observed for having conserved RNA structural elements (Rangan, Zheludev et al. 2020). The 12-nt insert, which does not exist in Spike region of SARS-CoV-1, also contains unusually high GC composition, increasing its likelihood to have a role on the RNA level as well as protein level. In this work, investigate the Spike gene on an RNA level. We compare the original SARS-CoV-2 viral strain with its closest relative SAR-CoV-1 for any sign of major long-range RNA-RNA interactions that involve a genomic segment on the *Spike* region. The impact of locally stable RNA structures on the long-range predictions are also investigated. Subsequently, we considered the population of evolving SARS-CoV-2 sequences available worldwide to further investigate the conservation of our inferred interactions.

## Materials and Methods

### Data

We used the SARS-CoV-2 (NC_045512) and SARS-CoV (NC_004718) reference sequences for identifying long-range RNA-RNA interactions in each of the viruses. For population-based sequence-covariance analyses, a set of 2’348’494 aligned full-length SARS-CoV-2 genome sequences were taken from the Nextstrain project (Hadfield, Megill et al. 2018) on December 9, 2021. The sequences were originally from the Global Initiative on Sharing All Influenza Data (GISAID) platform (Elbe and Buckland-Merrett 2017, Shu and McCauley 2017, Khare, Gurry et al. 2021) (https://www.gisaid.org/) and were subsequently filtered for high quality sequence (nextstrain.org, filename: filtered.fasta.xz). We further filtered the sequences for having no ambiguous nucleotides in desired locations which resulted in a total of 2’068’427 sequences. Supplementary Table 1 contains the corresponding GISAID accession numbers for the sequences. Finally, we performed down-sampling to around 10 percent of original size (206’745 sequences) due to computational complexity constraints.

### Predicting RNA-RNA interactions

Genome-wide RNA-RNA interaction between the Spike region (query) and the genomic RNA of SARS-CoV-2 (target) were predicted using IntaRNA (Busch, Richter et al. 2008, Wright, Georg et al. 2014, Mann, Wright et al. 2017, Raden, Ali et al. 2018). First, the Spike region was divided into smaller regions using a sliding window of length 500nt and overlap of 50nt. Each segment was then used as the query parameter by IntaRNA using search mode parameters (`--mode H --outNumber 5 --outOverlap Q`). The parameters allowed for extracting top 5 non-overlapping targets on the full genome that form thermodynamically favorable RNA-RNA base-pairing interactions with a region on the corresponding query segment. Targets that were at least 1000nt apart from their query counterparts were subsequently kept. A similar procedure was carried out on SARS-CoV-1.

Different components of the RNAstructure software package (Reuter and Mathews 2010) along with other tools were used for secondary structure predictions. Bimolecular RNA structural predictions were performed using bifold (Mathews, Burkard et al. 1999) default parameters. The bifold program takes two sequences as arguments and predicts the most energetically favorable bimolecular conformation. Minimum Free Energy structure (MFE) prediction was performed using both fold (Mathews 2004) and ViennaRNA software package (Lorenz, Bernhart et al. 2011) using default parameters. Individual base-pair probabilities are according to McCaskill’s partition function (McCaskill 1990, Mathews 2004). The Maximum Expected Accuracy (MEA) (Lu, Gloor et al. 2009, Reuter and Mathews 2010) and pseudoknot predictions were done using ProbKnot (Bellaousov and Mathews 2010, Zhang, Zhang et al. 2020) using default parameters.

### Compensatory Mutations Analysis of Long-range RNA-RNA interactions

Compensatory mutations within the multiple sequence alignments were investigated using the R-scape software package (Rivas, Clements et al. 2017, Rivas 2020, Rivas, Clements et al. 2020, Rivas and Eddy 2020), which analyzes covariation in nucleotide pairs in the population to infer possible compensatory mutations in an RNA base pair. If the consensus RNA secondary structure is not provided by the user, the software is also capable of predicting the consensus structure from the population of sequences using an implementation of the CaCoFold algorithm.

Compensatory (covarying) mutations for long-range RNA-RNA interactions were analyzed by retrieving the two sequence segments that constitute the desired RNA-RNA interaction for all downloaded SARS-CoV-2 sequences. Pairs of sequence segments were extended on each of their ends by 5nt (totaling 20nt) and concatenated. Then, the long-range RNA-RNA interacting structure was predicted by finding the consensus secondary structure within the population of sequences in the dataset using R-scape implementation of CaCoFold. The consensus structure was compared to bifold predictions for verification. Nucleotide pairs belonging to the consensus structure were then examined within the dataset for evidence of covariation using the built-in survival function that plots the distribution of base pairs with respect to their corresponding covariation scores.

### Results

Long-range RNA-RNA base-pairing interactions were predicted between the *Spike* region and the full genome for both SARS-CoV-1 and SARS-CoV-2 using IntaRNA software package (Figure 1). For each genome, the *Spike* region was extended 50nt on both directions. *Spike* sequence segments of length 500nt were analyzed separately for possible long-range interactions with their corresponding genomes (See Materials and Methods for details). We considered a maximum of five hits (the optimal interaction and another four sub-optimal interactions) for each analysis. Figure 1 shows the location of all the hits in both the genomes.

**Figure 1.**
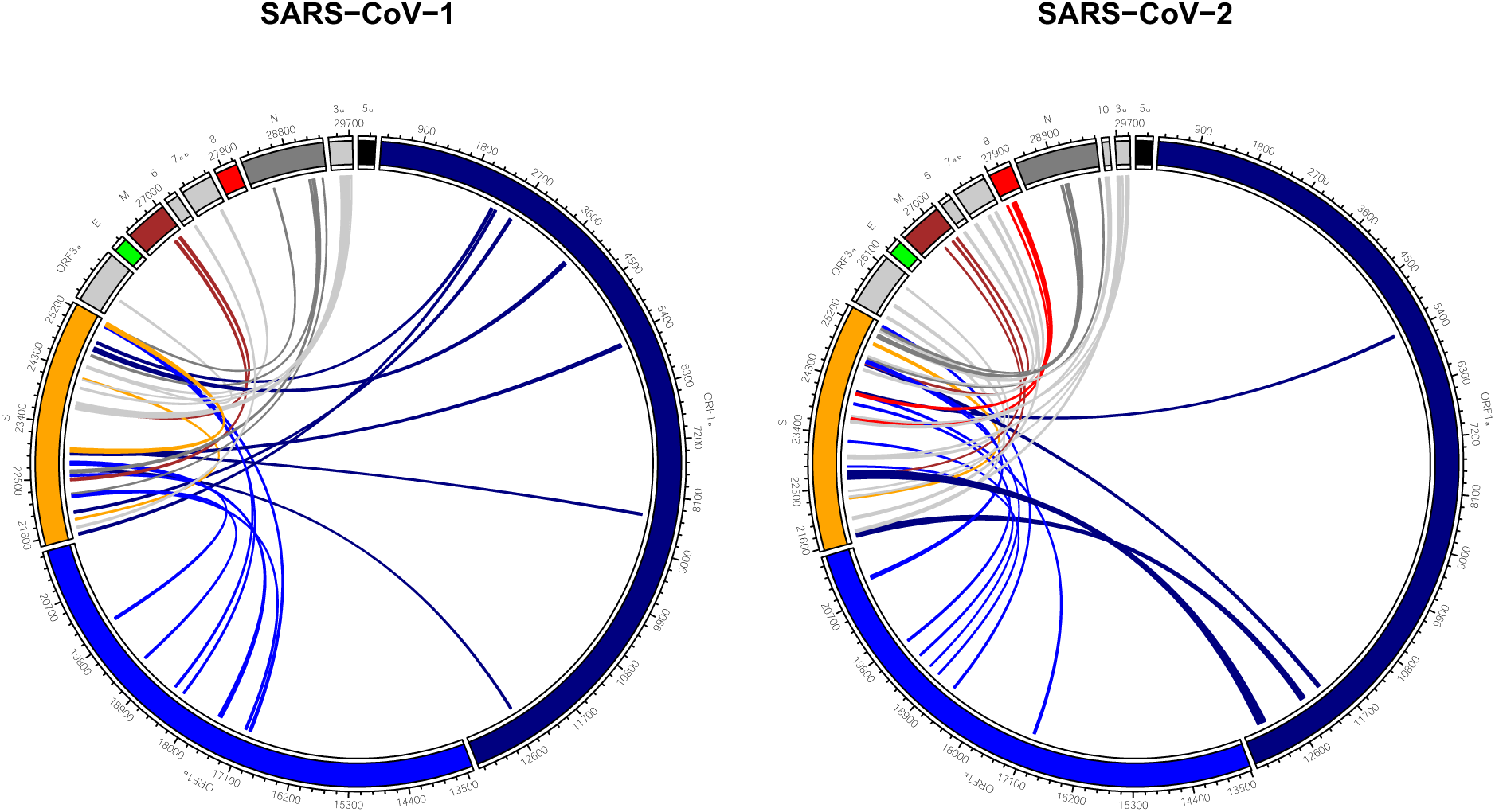
Predicted long-range RNA-RNA base-pairing interactions between *Spike* and the full genomic RNA. *Spike* sequence segments of length 500nt and overlap of 50nt were queried against the full genomes using IntaRNA software package. Each individual test resulted in at most five hits. All hits are summarized for both SARS-CoV-1 and SARS-CoV-2 (See Materials and Methods for details).

Long-range RNA-RNA predictions between *Spike* and the full genome are spread across almost all other genes for both SARS-CoV-1 and SARS-CoV-2 genomes. These interactions consisted of different thermodynamic stabilities and included interacting regions of as short as around 20nt. Supplementary Table 2 contains details about each hit. There were some major observations in our comparison. First, no interacting candidate was observed between the *Spike* and *E* genes for neither of the stains. Second, unlike SARS-CoV-2, a considerably long segment on SARS-CoV-1 *Spike* gene did not contain any prediction with the rest of the genome. In fact, the query segment of SARS-CoV-1 *Spike* (23’238 -23’737) contained only two hits, while other segments (on both strains) resulted at least four long-range predictions. The no-hit region corresponded to (23238-23698) on SARS-CoV-1, in specific. Finally, no prediction was observed between the SARS-CoV-1 *Spike* and *ORF8* regions, while this was not true for SARS-CoV-2. As we can see in Figure 1, there are multiple hits between *Spike* and *ORF8* for SARS-CoV-2.

There was a total of 69 long-range interactions across both strains. Table 1 summarizes the top quantile hits. The ranking of interactions was based on using their residual values against a generalized linear model that estimates interaction energy from interaction length. Model Akaike Information Criterion (AIC) was 365.76. (See Table 1 caption for Model details). Focusing only on SARS-CoV-2 hits, the top hit corresponds to the beginning of the *Spike* gene. In fact, the interaction overlaps with the upstream region of *Spike*. Interestingly, the second and third top hits are exactly adjacent to each other on the *Spike* region. Target regions shown in rank 3 and rank 7 are (24114-24157 *Spike*) and (24084-24114 *Spike*) and interact with their corresponding regions on Orf1a and Orf1b, respectively. Base-pair level interaction details for the top three interactions can be found in in Supplementary Figure 1.

**Table 1.** Top Quantile Predicted long-range RNA-RNA base-pairing interactions between the *Spike* region the full genome for both SARS-CoV-1 and SARS-CoV-2 using IntaRNA software package. See Materials and Methods for details. There was a total of 69 independent hits across both genomes. Complete results included as Supplementary Table 2. Column `SARS-CoV` denotes the strain. Column `TotalLength` denotes length of the interacting regions (query + target). Ranking is according to residual values against the generalized linear model where length of interaction was used to estimate interaction energy. Model Akaike Information Criterion (AIC) = 365.76. Length coefficient = -0.03190. Length was a significant factor in the model. (Pr(>|t|) for length = 0.00067. Median of residuals = -0.2287). 1-Quantile of residuals = -2.1536. SARS-CoV-2 hits are shown as bold.

| Rank | SARS-CoV | Hit Start | Hit End | Target Start | Target End | Total Length | Energy | Residual | Target Gene |
| --- | --- | --- | --- | --- | --- | --- | --- | --- | --- |
| 1 | 2 | **21639** | **21750** | **12261** | **12355** | **207** | **-26.29** | **-7.3397887** | **ORF1a** |
| 2 | 1 | 22604 | 22631 | 12507 | 12532 | 54 | -21.23 | -7.1602061 | ORF1a |
| 3 | 2 | **24114** | **24157** | **5367** | **5402** | **80** | **-19.93** | **-5.0308541** | **ORF1a** |
| 4 | 1 | 24396 | 24414 | 25582 | 25602 | 40 | -18.19 | -4.5667802 | ORF3a |
| 5 | 1 | 24841 | 24877 | 2247 | 2288 | 79 | -19.26 | -4.3927522 | ORF1a |
| 6 | 1 | 25198 | 25239 | 17014 | 17053 | 82 | -19.1 | -4.1370578 | ORF1b |
| 7 | 2 | **24084** | **24114** | **17012** | **17046** | **66** | **-18.4** | **-3.9474282** | **ORF1b** |
| 8 | 1 | 23698 | 23734 | 26957 | 27000 | 81 | -18.87 | -3.9389559 | M |
| 9 | 2 | **23271** | **23295** | **19401** | **19423** | **48** | **-17.4** | **-3.521595** | **ORF1b** |
| 10 | 2 | **22846** | **22862** | **18954** | **18970** | **34** | **-16.92** | **-3.4881691** | **ORF1b** |
| 11 | 2 | **23660** | **23703** | **28025** | **28060** | **80** | **-18.07** | **-3.1708541** | **8** |
| 12 | 2 | **22303** | **22337** | **24984** | **25023** | **75** | **-17.79** | **-3.0503449** | **S** |
| 13 | 2 | **24984** | **25023** | **22303** | **22337** | **75** | **-17.79** | **-3.0503449** | **S** |
| 14 | 1 | 21523 | 21559 | 2321 | 2354 | 71 | -17.53 | -2.9179375 | ORF1a |
| 15 | 2 | 25331 | 25358 | 18595 | 18620 | 54 | -16.79 | -2.7202061 | ORF1b |
| 16 | 1 | 24667 | 24677 | 24104 | 24114 | 22 | -15.37 | -2.320947 | S |
| 17 | 2 | **24648** | **24660** | **19153** | **19165** | **26** | **-15.44** | **-2.2633543** | **ORF1b** |
| 18 | 1 | 22530 | 22558 | 20039 | 20077 | 68 | -16.67 | -2.1536319 | ORF1b |

### *Spike*-*ORF8* RNA-RNA Interaction

The interaction between *Spike* and *ORF8* with the highest ranking appears as the 11^th^ top hit within a total of 66, under a generalized linear model that estimates interaction energy from sum of lengths of interacting sequences. It is also the 6^th^ top hit within SARS-CoV-2. Base-pairing interactions between SARS-CoV-2 *Spike* and *ORF8* are shown in Figure 2. Intervals (23660-23703 *Spike*) and (28025-28060 *ORF8*) consist of a total of 80nt and have a stabilizing energy of -18.07 kcal/Mol. Figure 2 shows the individual base pairs of the above hit, denoted here as *Spike-ORF8 interaction*. Pairs shown by ‘+’ symbol depict those that pair earlier than other base pairs (predictions according to IntaRNA).

**Figure 2.**
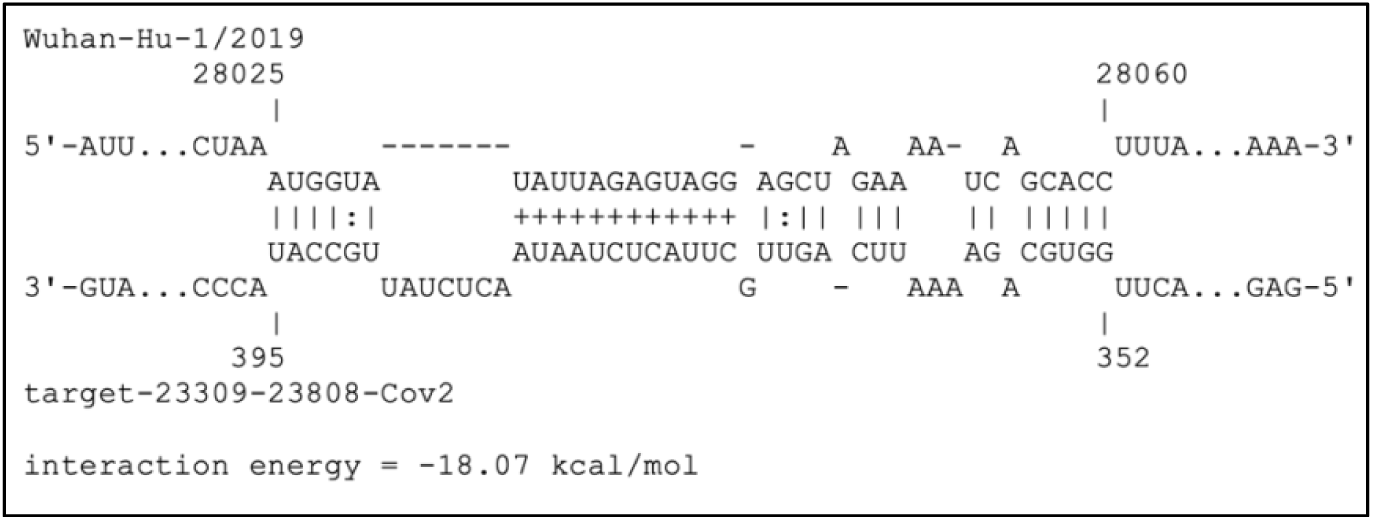
Long-range RNA-RNA interaction between *Spike* and *ORF8* regions of SARS-CoV-2 genome. Interacting intervals are (23660-23703 *Spike*) and (28025-28060 *ORF8*). Prediction done via IntraRNA software. Base pairs with ‘plus’ notation indicate an earlier binding than the other ones.

The predicted Spike-ORF8 interaction was analyzed for compensatory mutations. Sequence segments were extended 5nt to avoid unwanted base-pairing in the consensus structure prediction. Resulting intervals were (23655-23708 *Spike*) and (28020-28065 *ORF8*). A total of 206’745 sequence segments each corresponding to a particular viral strain were used for the analysis. Sequences were a down-sampled selection of nearly two million SARS-CoV-2 sequences (See Materials and Methods for detail). No significantly covarying mutations were detected by R-scape. Table 2 shows the coordinates of all base pairs for which variation was observed. Column power an output of the R-scape software, denotes the statistical power of substitutions.

**Table 2.** Coordinates of interacting base pairs between (23660-23703 *Spike*) and (28025-28060 *ORF8*) for which nucleotide variations were observed. Total number of sequences was 206745. Column `power` is an output of the R-scape software that is proportional to the statistical power of substitutions. Mutations in coordinates in red bold are shown Figure 3. The coordinates in black bold correspond to those pairing that fall in the region denoted by ‘+’ in Figure 2.

| *Spike* | *ORF8* | *Power* |
| --- | --- | --- |
| 23660 | 28060 | 0 |
| 23661 | 28059 | 0 |
| 23662 | 28058 | 0 |
| **23663** | **28057** | **0.08** |
| **23664** | **28056** | **0.11** |
| 23671 | 28050 | 0 |
| 23672 | 28049 | 0 |
| **23673** | **28048** | **0.39** |
| 23674 | 28046 | 0 |
| **23675** | **28045** | **0.05** |
| 23676 | 28044 | 0.04 |
| 23677 | 28043 | 0 |
| 23679 | 28042 | 0 |
| 23680 | 28041 | 0.01 |
| 23681 | 28040 | 0 |
| 23682 | 28039 | 0 |
| 23683 | 28038 | 0 |
| 23684 | 28037 | 0 |
| 23685 | 28036 | 0 |
| 23686 | 28035 | 0 |
| 23687 | 28034 | 0 |
| 23688 | 28033 | 0.01 |
| 23689 | 28032 | 0 |
| 23690 | 28031 | 0 |

Interestingly, comparing Figure 1 and Table 2, we can see that the base pairings that from earlier than others, shown with symbol ‘+’, happen to also have lower variation in the population of sequences than those base pairs that form afterwards. i.e., base pairs shown in black bold in Table 2 fall within the stem shown by symbol ‘+’ in Figure 2.

Figure 3 illustrates the individual pairing configurations within the RNA-RNA interaction. Results were according to the consensus structure prediction algorithm CaCoFold built in the R-scape software. All three structure predictions, IntaRNA (long-range), bifold (bimolecular folding), and CaCoFold (consensus structure) had consistent results. Four base pairs with highest number of mutations are shown. Nucleotide position with highest observed mutation was G28048U *ORF8*, with 36’366 occurrences in a total of 206’745 viral sequences. This mutation does not support the predicted interaction. Mutation C23664U *Spike* was observed 2207 times and was the second highest mutation observed. This mutation accommodates for the Spike-ORF8 interaction. Adjacent to this base pair, mutation G28045U *ORF8* with frequency 329 also accommodates for the interaction stability. The fourth most frequent mutation was C28045U *ORF8*. It was observed 329 times which also accommodated the predicted Spike-ORF8 interaction. The flanking sequences on both ends of interactions did not form any base pairing with each other as expected by IntaRNA results.

**Figure 3.**
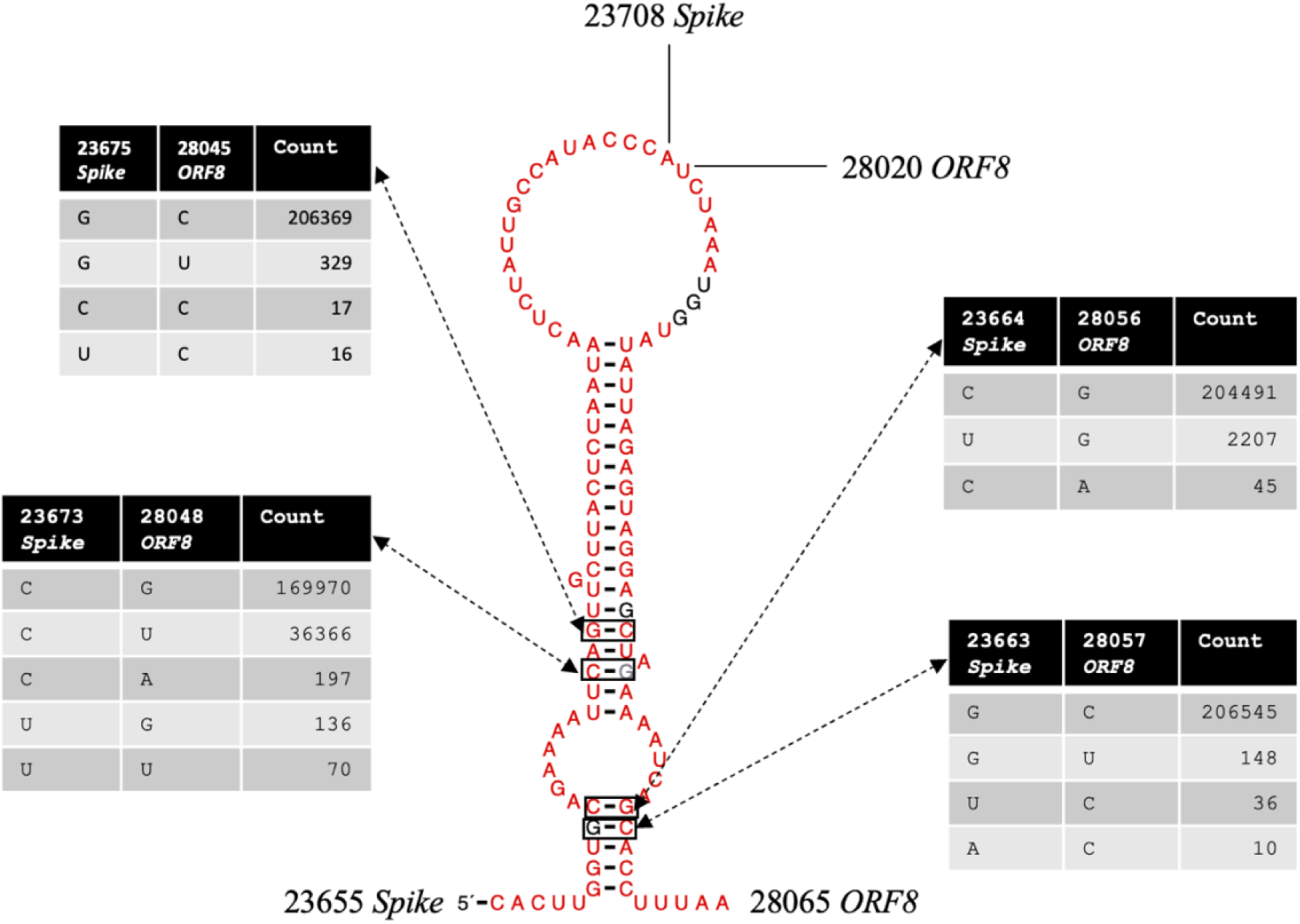
Consensus structure of the predicted RNA-RNA interaction (23660-23703 *Spike*) and (28025-28060 *ORF8*). Total number of sequences was 206’745. Number of mutations observed for four locations with highest power are shown.

### Local RNA Analysis in *Spike*

The local stability of RNA structure in the vicinity of the (23660-23703 *Spike*) was evaluated and compared to its SARS-CoV-1 counterpart. The original interval was extended by 100nt on both directions on the SARS-CoV-2 genome, resulting region (23560-23803 *Spike*). The region that aligned with the above selection on SARS-CoV-1 was selected for comparison, (23447-23650 *Spike*). Table 3 shows the pseudoknot structure prediction of the two regions. The SARS-CoV-2 segment contains five pseudoknots in its structure, while the SARS-CoV-1 counterpart contains none. This observation was consistent for a shorter interval selection as well, although the number and location of the pseudoknot would vary on the SARS-CoV-2 selected segment. Figure 4 shows the base pair probabilities for both SARS-CoV-2 (23560-23803 *Spike*) and its corresponding region in SARS-CoV-1 (23447-23650 *Spike*). As we can see, there are major differences in the base-pairing probability patterns between the two sequences. The line in bold shows the approximate location of the Spike-ORF8 interaction. As we can see this location seems to contain many bases pairs that can form local base pairs. The corresponding location on SARS-CoV-1, for which no long-range interaction with *ORF8* was observed, seems to have relatively less locally stable bases pairs (comparing red base pairs between Figure 4A and Figure 4B). This observation was also true for another arbitrary selection of sequence segments. Overall, region of *Spike* that is predicted to base pair with *ORF8*, also tends to form a local structure which seems to be mutually exclusive from the *ORF8* interaction. In addition, other structural signatures such as pseudoknots was observed in the vicinity of the mentioned region, while no sign of pseudoknot predictions was seen in that of SARS-CoV-1.

**Table 3.**
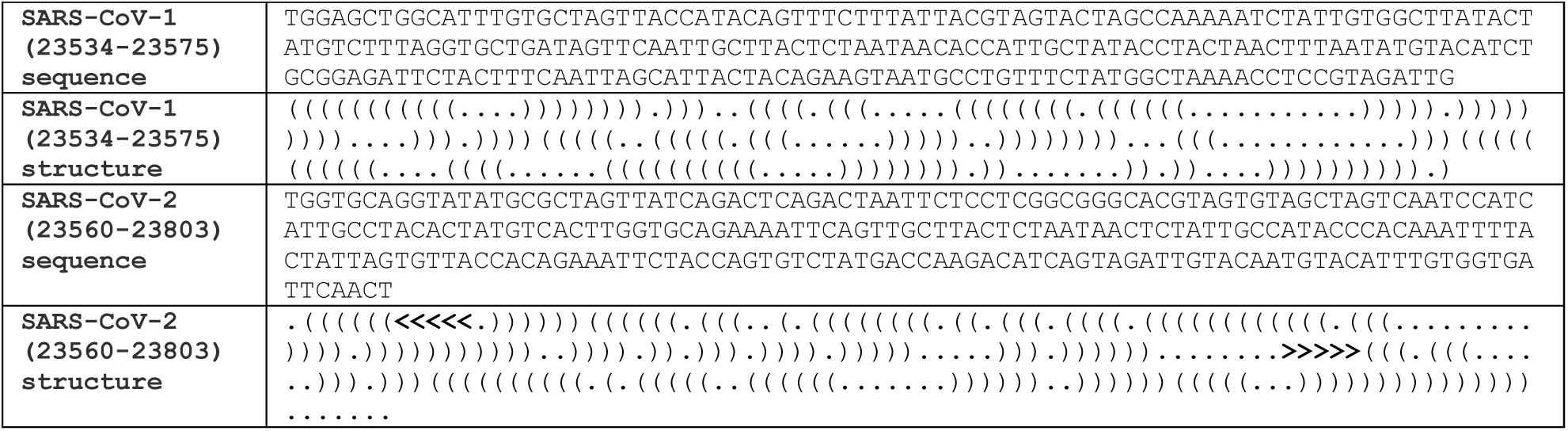
Pseudoknot structure prediction of aligned segments of *Spike* using Probknot. Sequences for SARS-CoV-1 (23447-23650 *Spike*) and SARS-CoV-2 (23560-23803 *Spike*) are shown along with their corresponding predictions.

**Figure 4.**
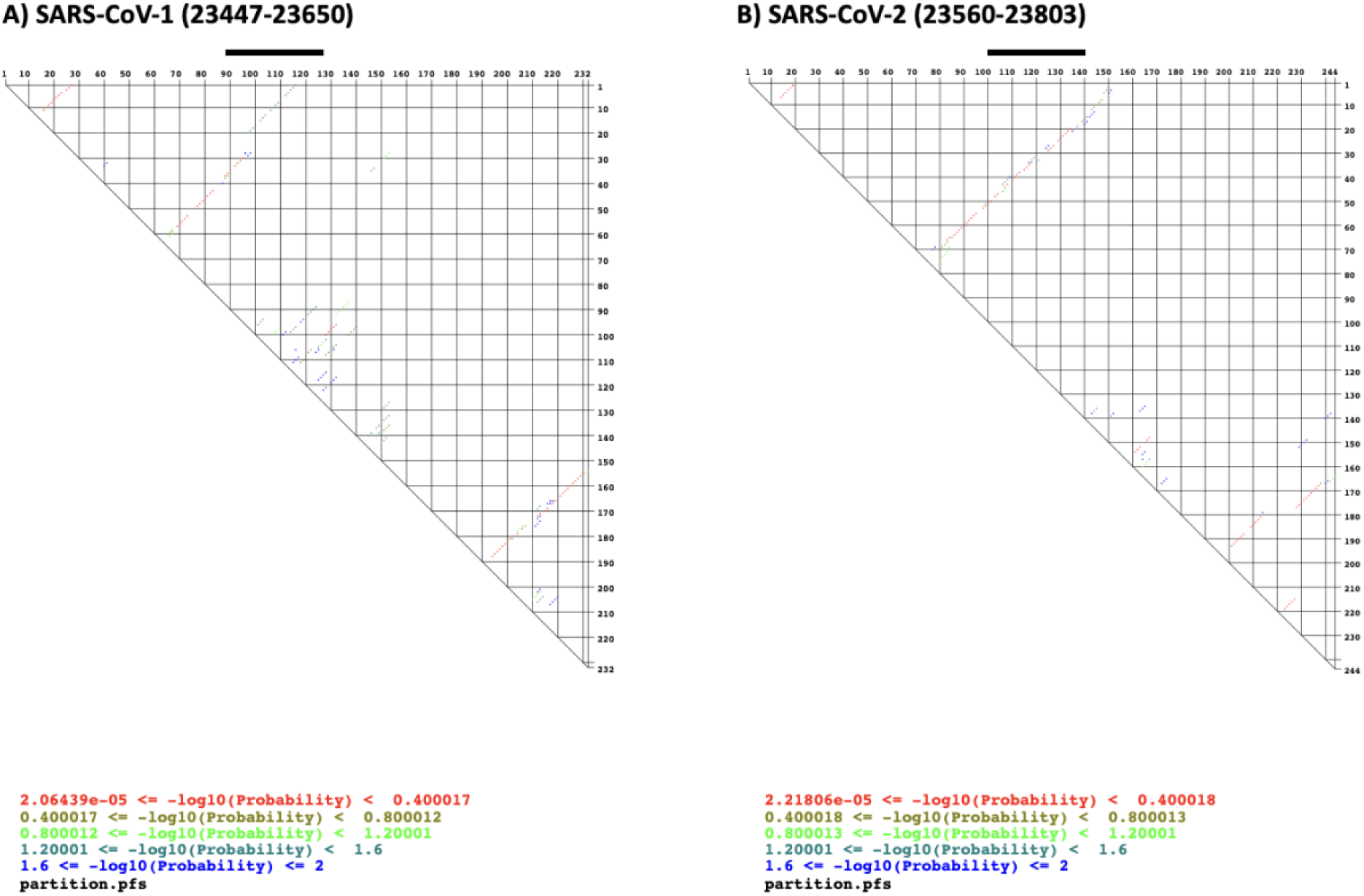
Base pair probabilities for aligned segments of *Spike* using the partition functions. A) SARS-CoV-1 (23447-23650 *Spike*) B) SARS-CoV-2 (23560-23803 *Spike*). The Spike/ORF8 region (23660-23703), shown as black line in (B) was extended on both directions by 100nt. The corresponding region, SARS-CoV-1 (23534-23575 *Spike*), is shown in bold in (A).

## Discussion

The Spike region of SARS-CoV-2 RNA was investigated for novel genomic long-range RNA-RNA interaction. *In-silico* predictions were performed on the reference sequence and compared to those for the reference sequence of SARS-CoV that was responsible for the 2002-2003 outbreak. The predictions were inclusive and made in favor of more sub-optimal but diverse results. They provide a collection of top non-overlapping candidate regions on the reference sequences that can potentially form thermodynamically favorable RNA-RNA base pairing with a sub-region on their corresponding *Spike* (Figure 1). We found RNA structural differences between corresponding regions in SARS-CoV-2 and SARS-CoV.

Top interacting regions were ranked according to their relative thermodynamic stabilities with regards to total length of interaction, using a generalized linear model. Table 2 shows the top quantile of results (See Supplementary Table 2 for full results). Some of the predictions are as follows. Strongest interactions that occurred on the SARS-CoV-2 genomic RNA were between *Spike* and *Orf1ab*. A region in the beginning of SARS-CoV-2 Spike (21639-21750) formed an interaction with a region on *Orf1ab* (12261-12355) with a predicted free energy of -26.29 kcal/Mol, highest amongst both viruses. Details about base pairing interactions of the top three hits is presented in Supplementary Figure 1. The second and third strongest predictions on SARS-CoV-2 also occurred on Orf1ab but formed continuous region on the side of *Spike*. Regions (24114-24157 *Spike*) and (24084-24114 *Spike*) intersect at position 24114 but interact with distant regions on *Orf1ab*, namely the beginning (5367-5402) and the middle (17012-17046), respectively. This observation was unique, since interactions were allowed to overlap on *Spike* by the IntaRNA software, but they only have one nucleotide overlap on *Spike* (See Table 2, rows 3 and 7 for details). Some of the other interesting observations were the fact that a significantly long region of SARS-CoV-1 *Spike* (23238-23698), around 460nt, did not form any long-range RNA-RNA predictions with any part of the genome, despite the software’s flexibility to allow for sub-optimal hits. This lack of predictions was not observed in SARS-CoV-2 *Spike*.

Most genes and annotated regions contained several interacting regions with the Spike gene in both reference genomes SARS-CoV-1 and SARS-CoV-2 (comparing Figure 1A and Figure 1B). The ranking of strength of base-pairings, however, were dramatically different between corresponding genes. For instance, the strongest ranked interaction between *Spike* and *M* in SARS-CoV-1 was 8^th^ while this number dropped to 40 for SARS-CoV-2 (See Supplementary Table 2). The only gene that did not contain any predictions was the *E* gene. No thermodynamically stable interacting candidate was observed on neither of the strains of SARS-CoV-1 and SARS-CoV-2 reference genomes. In the case of *ORF8*, however, SARS-CoV-2 contained a few regions that can potentially form long-range RNA-RNA interactions with the *Spike* region on different locations (Figure 1B, red links), while SARS-CoV-1 didn’t contain any. The ranking of the highest observed interaction stability fell within the first quantile of results (ranking 11). Regions (23660-23703 *Spike*) and (28025-28060 *ORF8*) formed an RNA-RNA interaction with free energy of -18.07 kcal/Mol. While the other results were interesting and worth further investigation, our focus was further analysis of the above Spike-ORF8 interaction, due to the strong gene-based observed contrast between the SARS-CoV-1 and SARS-CoV-2.

The population of SARS-CoV-2 sequences were analyzed for signs of sequence co-variation that might validate the above S-ORF8 RNA-RNA base-pairing interaction. From amongst the nearly 20 million sequences, 206’745 (roughly 10%) were randomly selected for the analysis, due to limitations in computational complexity. The aligned SARS-CoV-2 sequences were investigated for compensatory mutations that might occur within and between *Spike*-*ORF8* binding location. Although not any significantly covarying mutations were observed, the positions of polymorphisms were in support of the *in-silico* results. The *in-silico* predictions made on the reference sequences are based on base-pairs that from first which are more critical for the predicted interaction to stabilize. Interestingly those regions that pair first, are also the ones that were observed to tolerate less mutations (Figure 2; base pairs shown with ‘+’, Table 2; rows shown in bold). The lower variance in the more critical base pairs suggests further evidence for the Spike-ORF8 RNA-RNA interaction.

Observed mutations within the interacting region, however, had conflicting implications, with some such as C28045U *ORF8*, G28048U *ORF8*, C23664U *Spike* being in favor of the interactions and some such as G28048U not accommodating for base pairing (Figure 3). Being a synonymous mutation, C28045U has been previously identified as one of the polymorphic positions of *ORF8* (Pereira 2020). In the mentioned work, in the local RNA secondary structure prediction of *ORF8*, C28045U is unpaired, while in the predicted long-range RNA-RNA interaction with *Spike*, it pairs with G23675. The C28045U variation, hence, is suggestive of the long-range Spike-ORF8 interaction. Further investigation on the above set of mutations along the evolutionary trajectory of the virus is needed for a more comprehensive conclusion about their possible RNA structural roles. In addition, since the data was filtered and aligned for having no long inspersions, deletions, or ambiguous nucleotides, certain meaningful sequence variations might not have been accounted for in the analysis.

Local RNA structure analyses on the *Spike* region suggests an increase in locally stable RNA structures in the vicinity of the Spike-ORF8 interaction. *In-silico* local structure prediction of surrounding regions of the *Spike* target predicts the formation of pseudoknots in the SARS-CoV-2 region but not in the corresponding SARS-CoV-1 counterpart (Table 3). There is also a conserved RNA stem-loop, namely S1, which has been previously found in SAR-CoV sequences (Rangan, Zheludev et al. 2020). This stem is roughly 30nt upstream of the Spike-ORF8 interaction and its stability was confirmed by different *in-silico* programs in both SARS-CoV-1 and SAR-CoV-2 sequences. Immediately upstream of the conserved stem, there is the high-GC content 12-nt insert in the *Spike* region (23’603-23’614), which is present in SARS-CoV-2 but absent in SARS-CoV-1. The insert is roughly 50nt upstream of the predicted Spike-ORF8 interaction. Given the above comparisons to SARS-CoV-1, it seems that this region of *Spike* is undergoing local RNA structural changes as well as well as having affinity to form a long-range interaction with ORF8.

Locally stable RNA base pairs and the long-range Spike-ORF8 base-pairing interactions are mutually exclusive. Base pair probability distributions of corresponding regions on *Spike* in both SARS-CoV-1 and SARS-CoV-2 reveal that the same nucleotides that can pair with ORF8, are also likely to form local base pairs within *Spike* (Figure 3, red base pairs within the bold line). Ironically, the corresponding region on SARS-CoV-1, for which there were no signs of long-range interaction with ORF8, is observed to have less deterministic local base-pairing probabilities (Comparing Figure 4A and 4B, range indicated by bold lines). One possibility is that a complex RNA structure may be emerging within the specified region of *Spike* in SARS-CoV-2 that can form RNA-RNA interaction with ORF8, at certain times can avoid the interaction at others. Whether the predicted long-range Spike-ORF8 interaction is in competition or cooperation with other local elements of *Spike* such as the 12-nt polybasic insert in SARS-CoV-2, is subject to speculation about *in-vivo* conformational specifics.

The predicted *Spike*-*ORF8* long-range RNA-RNA interaction can potentially impact template switch during negative strand synthesis. Template switch in Beta-coronaviruses might occur if the TSR element downstream of the 5’UTR is in proximity of the TSR element immediately upstream of a viral gene. Such complex genomic conformation may involve other RNA-RNA as mediators. The dE-pE (Figure 2 of (Sola, Almazán et al. 2015)) act as such mediator RNA binding locations to facilitate a discontinuous negative strand synthesis of the viral genome, leading to N-gene sub-genomic RNA. The coronavirus nucleocapsid (N) is known to be a structural protein that forms complexes with genomic RNA, interacts with the viral membrane protein during virion assembly and plays a critical role in enhancing the efficiency of virus transcription and assembly (Sola, Almazán et al. 2015). The predicted Spike-ORF8 interaction here is 200nt upstream of the N-gene TSR (Pereira 2020). Although high-order RNA-RNA interactions needed for template switch can be more complex and may involve the 5’UTR as well, the predicted Spike-ORF8 interaction could be acting as an additional mediator step to bring the TRS elements of 5’UTR and the coronavirus *N*-gene closer to each other. It could be speculated that the Spike-ORF8 interaction is taking part in regulating sub-genomic RNA production. Since the first gene downstream of Spike-ORF8 interaction happens to be the *N*-gene, the binding location might be affecting the *N*-gene sub-genomic RNA production.

Amongst coronaviruses, *ORF8* is a rapidly evolving hypervariable gene that undergoes deletions to possibly adapt to human host (Gong, Tsao et al. 2020, Pereira 2020, Su, Anderson et al. 2020, Zinzula 2021). It has also been previously observed that patients infected with SARS-CoV-2 variants with a 382-nucleotide deletion (Δ382) in *ORF8* had milder symptoms (Young, Fong et al. 2020). In addition, *ORF8* contains RNA structural features (Pereira 2020). While this observation may very well be due to impact of absence of the translated protein, *ORF8* RNA structural characteristics of the genome may also play a role in the viral life cycle, making long-range RNA-RNA prediction with *Spike* a less remote possibility. A comprehensive exploration of predicted Spike-ORF8 interaction in other SARS-CoV-2 variants and evaluating corresponding sub-genomic RNA production rates of these variants may lead to further clues about the predicted long-range Spike-ORF8 RNA-RNA interaction, which can be rewarding for therapeutic purposes.

