## Supplementary Figure 1 for "Evidence for a long-range RNA-RNA interaction between *ORF8* and *Spike* of SARS-CoV-2"

Hit 1 (Ranking 1)

```
Wuhan-Hu-1/2019
         12261                                                                                                                   12355
             |                                                                                                                   |
5'-AUU...CAAC    -           U-  CU-  U    -  A   CCCA      U    A  CUA           AAG---    C  A  U    ---------------    C       ACAA...AAA-3'
             GUAA GUUGGAAAAGA  GG   GA CAAG CU UGA    AAUGUA AAAC GG   GAUCUGAGGAC      AGGG AA AG UACU               AGUG UAUGCAG
             |||| +++++++++++  ||   || |||| |: |||    |||||| |||| |:   +++++++++++      |||| || |: :||:               |||| +++++++
             CAUU UAACCUUUUCU  CC   CU GUUC GG ACU    UUACAU UUUG CU   CUAGACUUUUG      UCCC UU UU GUGG               UCAC AUACGUC
3'-AAA...GGUU    G           UU  AUU  U    A  -   CAAC      -    A  C--           AAACAG    A  A  U    UGCACACUUUCUUAA    -       CCCC...AAC-5'
             |                                                                                                                   |
           238                                                                                                                   127
target-21513-22012-Cov2

interaction energy = -26.29 kcal/mol
```

Hits 2 & 3 (Rankings 3 & 7)

```
Wuhan-Hu-1/2019
          5367                                           5402
             |                                           |
5'-AUU...UACA         C         U-----   ----             UUAU...AAA-3'
             GAGCAAGGG UGGUGAAGC      GCU    AACUUUUGUGCAC
             |++++++++ |||::||:|      ||:    +++++++++++++
             CUCGUUUCC ACCGUUUUG      CGG    UUGAAAACACGUG
3'-GUU...GACA         -         UCAUUC   CAAU             UUUA...AAC-5'
             |                                           |
           400                                           357
target-23758-24257-Cov2

interaction energy = -19.93 kcal/mol


Wuhan-Hu-1/2019
         17012                                  17046
             |                                  |
5'-AUU...AUCU        U         AUGUU    -        AAGG...AAA-3'
             CAGAUGAG UUUCUAGCA     GCAA AUUAUCAA
             ++++++++ +++++++++     |||| ++++++++
             GUUUACUC AGAGAUCGU     CGUU UAGUGGUU
3'-GUU...ACGU        C         -----    A        CCGU...AAC-5'
             |                                  |
           357                                  327
target-23758-24257-Cov2

interaction energy = -18.4 kcal/mol
```
